# Rapid evolution promotes fluctuation-dependent species coexistence

**DOI:** 10.1101/2020.08.05.239053

**Authors:** Masato Yamamichi, Andrew D. Letten

## Abstract

Recent studies have demonstrated that rapid contemporary evolution can play a significant role in regulating population dynamics on ecological timescales. Here we identify a previously unrecognized mode by which rapid evolution can promote species coexistence via temporal fluctuations and a trade-off between competitive ability and the speed of adaptive evolution. We show that this interaction between rapid evolution and temporal fluctuations not only increases the range of coexistence conditions under a gleaner-opportunist trade-off (i.e., low minimum resource requirement [*R*\*] vs. high maximum growth rate), but also yields stable coexistence in the absence of a classical gleaner-opportunist trade-off. Given the propensity for both oscillatory dynamics and divergent rates of adaptation (including rapid evolution and phenotypic plasticity) in the real world, we argue that this expansion of fluctuation-dependent coexistence theory provides an important overlooked solution to the so-called ‘paradox of the plankton’.

## Introduction

A wealth of empirical evidence has accumulated over the last few decades indicating that rapid contemporary evolution on ecological timescales (Hendry 2016) can be essential for understanding population dynamics (Yoshida *et al.* 2003; Bell 2017). More recently, evidence has begun to emerge that rapid evolution can also be a significant driver of community dynamics amongst competing species, and in particular play an important role in regulating species coexistence (Lankau 2011; Vasseur *et al.* 2011; Mougi 2013; Hiltunen *et al.* 2017; Wittmann & Fukami 2018; Hart *et al.* 2019; Germain *et al.* 2020; van Velzen 2020; Yamamichi *et al.* 2020). To date, however, rapid evolution has only been shown to promote species coexistence when there is either a trade-off between traits optimal for intraspecific and interspecific competition (Lankau 2011; Vasseur *et al.* 2011; Mougi 2013; Wittmann & Fukami 2018; Yamamichi *et al.* 2020) or fine-tuning of prey defenses and predator foraging efforts (Kondoh 2003; van Velzen 2020). Here we provide a previously unrecognized pathway for rapid evolution to promote coexistence via temporal fluctuations: differences in the rate of adaptation to fluctuating environments. Specifically, using model simulations, we show that rapidly evolving and non-evolving consumers can stably coexist even when they compete for a single resource.

Understanding how species coexist in nature has been a central topic in community ecology over the last century (Hutchinson 1961; Wilson 1990). G. E. Hutchinson (1961) famously referred to the problem as “the paradox of the plankton” on account of the seemingly impossible challenge of explaining the extraordinary diversity of oceanic plankton (Hutchinson 1961). Independent of emerging interest into rapid evolution, a growing body of coexistence research has suggested that temporal fluctuations can promote diversity via two broad classes of mechanisms, the temporal storage effect and relative nonlinearity (Chesson 1994, 2000; Yuan & Chesson 2015; Barabás *et al.* 2018; Ellner *et al.* 2019). Although greater attention has been given to the temporal storage effect (Adler *et al.* 2006; Angert *et al.* 2009; Letten *et al.* 2018; Hallett *et al.* 2019; Zepeda & Martorell 2019), recent studies have shown that relative nonlinearity can be more important than previously thought for promoting coexistence (Letten *et al.* 2018; Hallett *et al.* 2019; Zepeda & Martorell 2019). In this paper, we introduce rapid evolution as a previously neglected driver of relative nonlinearity.

Relative nonlinearity promotes coexistence when one species benefits from fluctuations in the intensity of competition (fluctuation-assisted) while the other species dominates in the absence of fluctuations (fluctuation-impeded). In addition to their responses to fluctuations, for the two competing species to coexist, they need different impacts on fluctuations. If the fluctuation-assisted species reduces the amplitude of fluctuations (by “consuming” the variance in the intensity of competition (Levins 1979)) while the fluctuation-impeded species increases it, each species produces a favorable environment for the competitor, resulting in negative frequency-dependence in community dynamics (Hsu *et al.* 1978; Armstrong & McGehee 1980) (Fig. 1a-b).

**Figure 1.**
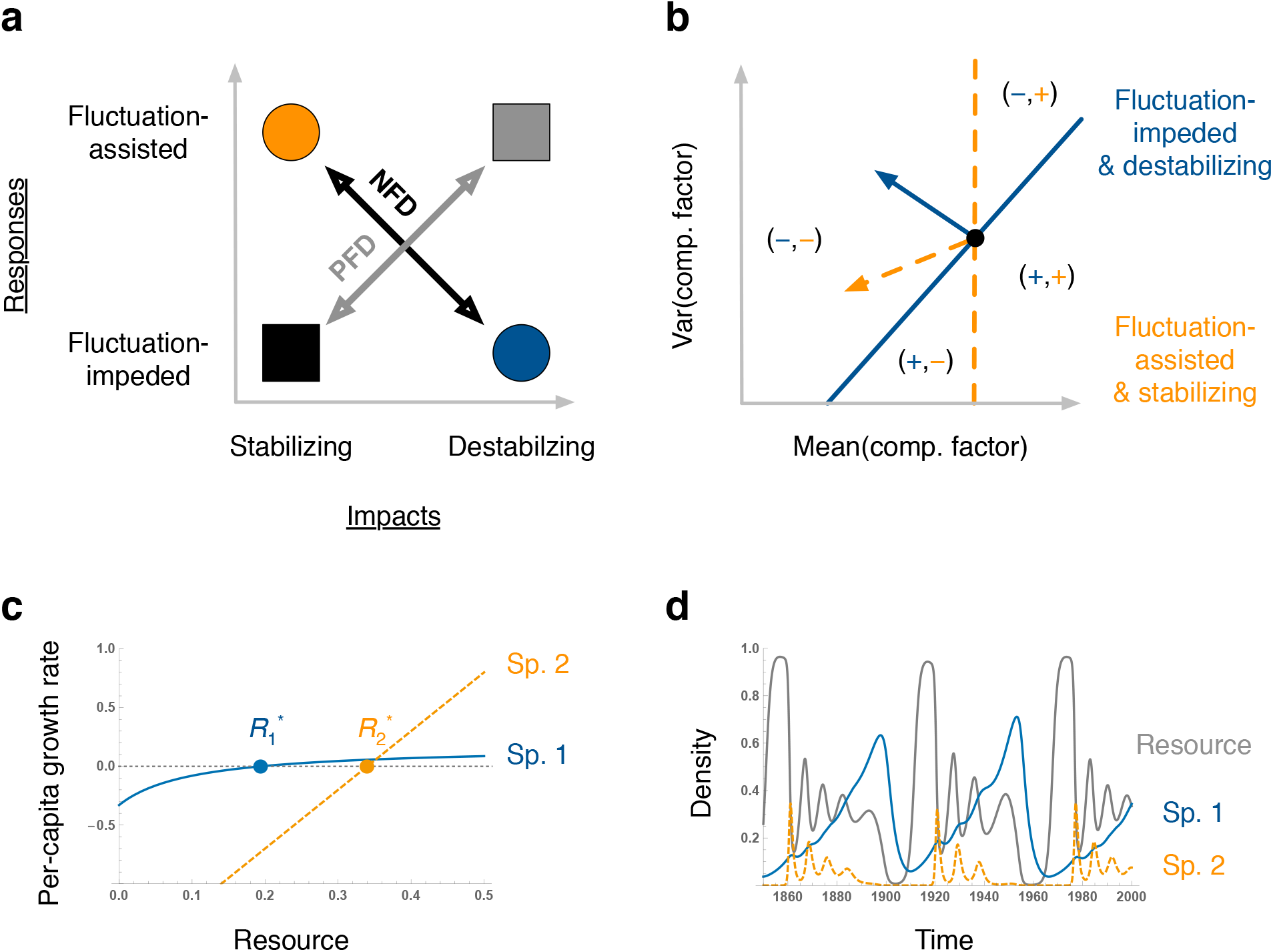
Species coexistence by relative nonlinearity. **a**, Relative nonlinearity promotes coexistence with negative frequency-dependence (NFD) when one species stabilizes population cycles but benefits from fluctuating environments in competition (fluctuation-assisted), while the other species destabilizes dynamics but benefits from stable environments in competition (fluctuation-impeded). Hence, we need to consider two aspects of the ecological niche: responses and impacts (the Grinnellian and Eltonian niches, respectively: note that with a feedback cycle, the ideas of response and effect are complicated since the two terms rely heavily on perspective) (Chase & Leibold 2003). Priority effects due to positive frequency-dependence (PFD) can arise when the fluctuation-impeded species stabilizes fluctuations while the fluctuation-assisted species drive fluctuations (Ke & Letten 2018). **b**, The concept of relative nonlinearity can also be expressed in terms of zero net growth isoclines (ZNGI) (Chase & Leibold 2003). The fluctuation-assisted (fluctuation-impeded) species has higher growth rates when the variance in the competitive factor (e.g., resource) is high (low), but when abundant reduces (increases) the variance as shown by the impact vector. **c**-**d**, Coexistence of species with linear and saturating functional responses (Abrams *et al.* 2003). Species 1 with a saturating functional response (blue line) has a smaller *R*\* and causes limit cycles with the resource (grey line), which is disadvantageous due to nonlinear averaging. Species 2 with a linear functional response (the dashed orange line) has a larger *R*\* and stabilizes dynamics. Parameter values are *a*_1_ = *a*_2_ = 5, *b*_1_ = 10, *b*_2_ = 0, *d*_1_ = 0.33, and *d*_2_ = 1.7.

As the name suggests, two species may coexist via relative nonlinearity when consumer population growth responses to a limiting resource differ in their degree of nonlinearity (Fig. 1c-d) (Hsu *et al.* 1978; Armstrong & McGehee 1980; Abrams & Holt 2002; Abrams *et al.* 2003; Xiao & Fussmann 2013). The differential responses to, and impacts on, fluctuations are most conspicuous when competing consumers exhibit intermittent cycles (Fig. 1d), where the linear, competitively inferior species at equilibrium (with larger *R*\*), increases in abundance when the system oscillates but simultaneously dampens the consumer-resource cycles induced by the competitively superior species (with smaller *R*\*). In contrast, the nonlinear, competitively superior species generates fluctuations but increases in abundance when the system is stable (Fig. 1d). The species with the lowest *R*\* value would outcompete its competitor under a stable equilibrium, but the linear species benefits from fluctuating environments in competition due to nonlinear averaging of the low *R*\* species’ per capita growth response (Jensen’s inequality). The situation in Fig. 1c is called a gleaner-opportunist trade-off, where species 1 has the lower *R*\* whereas species 2 has the higher maximum growth rate (Grover 1990). Because this criterion is sometimes perceived as being highly restrictive, relative nonlinearity has typically been assumed to play a minor role in real systems (Adler *et al.* 2006; Angert *et al.* 2009; Letten *et al.* 2018; Hallett *et al.* 2019; Zepeda & Martorell 2019). By demonstrating how rapid evolution can relax these restrictions and significantly broaden opportunities for coexistence even without the classical gleaner-opportunist trade-off, we argue that relative nonlinearity may operate much more widely in nature than previously thought.

## Methods

The basis for the mechanism presented here is that rapid evolution can enable populations to adapt to fluctuating environments (response) and to stabilize temporal fluctuations (impact) (Cortez & Patel 2017). Thus, even when two species with fixed functional responses cannot coexist, they may have different responses to, and impacts on, temporal fluctuations, facilitating coexistence. We consider a model of two consumers with saturating consumer functional responses competing for a single logistically growing resource (Rosenzweig & MacArthur 1963; Hsu *et al.* 1978; Xiao & Fussmann 2013). In our model, the dynamics of the two consumers, *N*_1_ and *N*_2_, competing for a single resource, *R*, with quantitative traits, *x*_1_ and *x*_2_ is represented by

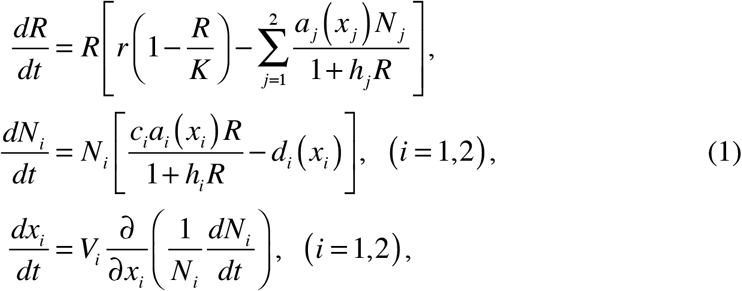

where *r* is the intrinsic growth rate, *K* is the carrying capacity, *a_i_* is the parameter that increases the consumption rate as a whole (including the slope and maximum value), *b_i_* is the parameter that reduces the maximum consumption rate, *c_i_* is the conversion efficiency, *d_i_* is the consumer mortality, and *V_i_* is additive genetic variance of the trait *x_i_* (*i* = 1, 2).

We assume a trade-off between parameters for the consumption rate, *a_i_*, and mortality rate, *d_i_*, such that increasing the consumption rate increase mortality rate in an accelerating manner (*a_i_*(*x_i_*) = *α_i_x_i_*, *d_i_*(*x_i_*) = *δ*_0*i*_ + *δ*_1*i*_*x*_*i*_ + *δ*_2*i*_*x_i_*^2^). This occurs due to, for example, a foraging-predation risk trade-off (Verdolin 2006) where predators have Holling type III functional responses. In addition, we assume the trait changes along the fitness gradient (Abrams 2001; Cortez & Patel 2017). Thus, trait dynamics can be represented by

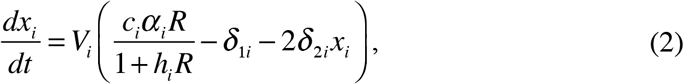

and the optimal trait at a certain resource level is obtained by solving *dx_i_*/*dt* = 0, and it is given by *x_i_* = (*c_i_α_i_R*/(1 + *b_i_R*) – *δ*_1*i*_)/(2*δ*_2*i*_). There is stabilizing (instead of disruptive) selection toward the optimal trait as *∂*(*dx_i_*/*dt*)/*∂x_i_* = −2*V_i_δ*_2*i*_ < 0. This indicates that the evolving consumer invests more in resource uptake when resources are abundant, whereas it tries to reduce its mortality rate when resources are scarce. Furthermore, by nondimensionalization, we can assume that *r* = *K* = *c_i_* = 1 without loss of generality (Appendix S1).

As the resource exhibits logistic growth and consumers exploit the resource with a Holling type II functional response, the system corresponds to the classical Rosenzweig-MacArthur model when there is a single non-evolving consumer species (Rosenzweig & MacArthur 1963). In the absence of fluctuations, the consumer with the smaller resource requirement (i.e., smaller *R*\*)reduces the resource level to the point 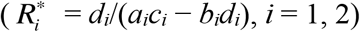 where its competitor with a larger *R*\* cannot persist and is excluded. On the other hand, in the presence of endogenous limit cycles (e.g., with larger *K* values) the two consumers may coexist via relative nonlinearity. Given Holling type II (concave) functional responses, population growth rates of the two species are lowered by fluctuations, but the species with lower *R*\* is more impeded by fluctuations in the intensity in competition, thereby promoting coexistence (Hsu *et al.* 1978; Xiao & Fussmann 2013).

## Results

When we assume that a competitively superior species (with smaller *R*\* at equilibrium) does not evolve, whereas an evolving, competitively inferior (larger *R*\*) species with the optimum trait either grows faster than the competitively superior species when the resource is abundant (Fig. 2a) or scarce (Fig. 2b), coexistence is possible. Notably, our parameter settings do not satisfy necessary conditions for coexistence via the classical gleaner-opportunist trade-off identified in previous studies (Appendix S1, Fig. S1) (Hsu *et al.* 1978; Xiao & Fussmann 2013) for any fixed trait values. Thus, when the speed of evolution in the species that is competitively inferior at equilibrium is slow, competitive exclusion ensues (Fig. S2). However, when evolution is faster, coexistence is possible (Fig. 2). This occurs because relative to the non-evolving species, the rapidly evolving species benefits from resource fluctuations, but at the same time rapid evolution dampens the consumer-resource cycles (Appendix S2, Fig. S3) induced by the non-evolving superior competitor, as can be clearly seen when the system cycles intermittently (Fig. 2f). Therefore, rapid evolution works as a compensation mechanism for the species that would otherwise be competitively inferior in a non-fluctuating system.

**Figure 2.**
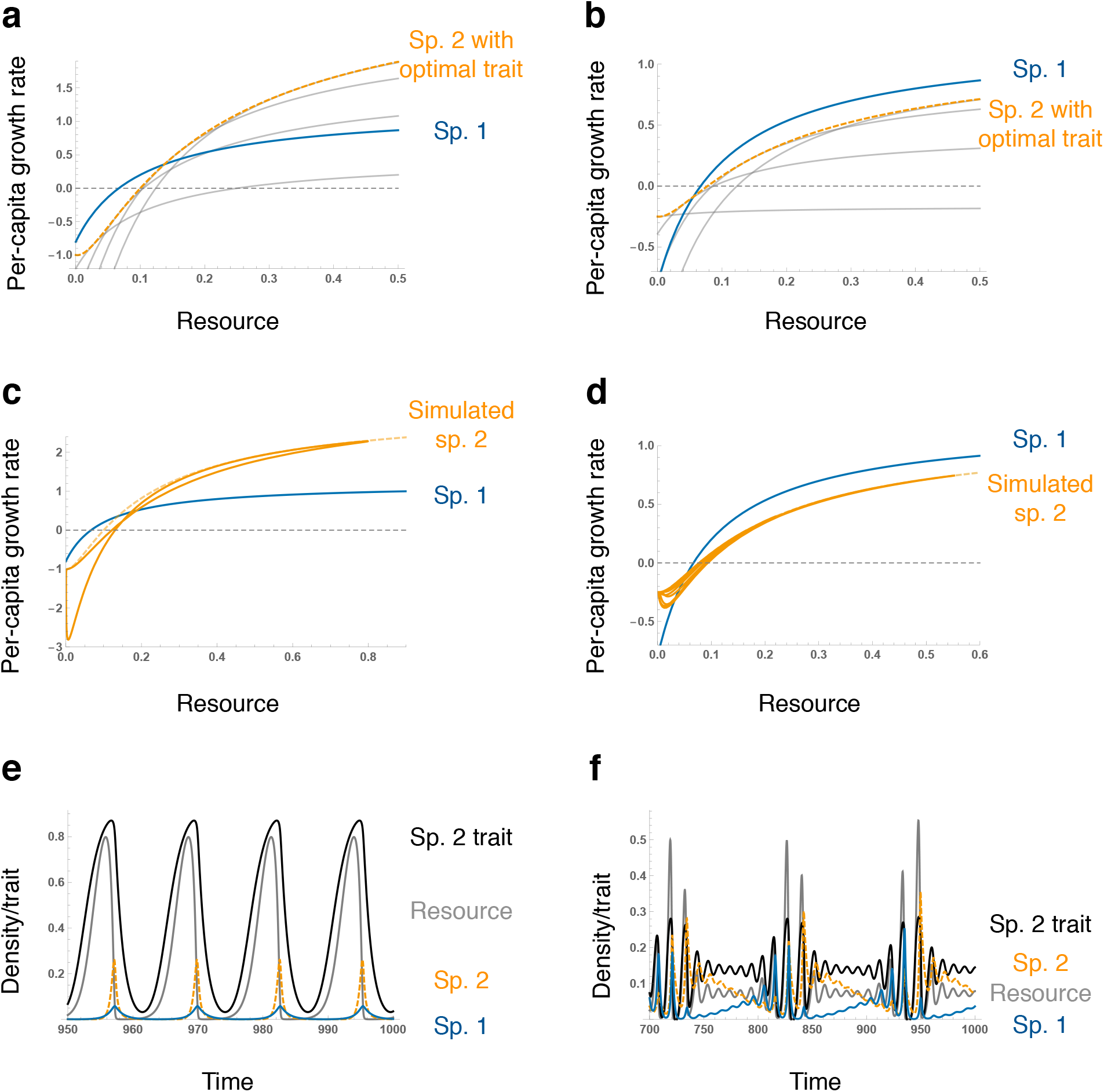
Rapid evolution of a competitively inferior (high *R*\*) species promotes coexistence by relative nonlinearity. **a-b**, The per-capita growth rate of species 2 with the optimal trait (orange dashed line) is larger than that of species 1 (solid blue line) when the resource is abundant (**a**) or scarce (**b**). When the trait value is fixed (gray lines), species 2 is always competitively inferior (Fig. S2). **c-d**, The realized per-capita growth rate of species 2 (i.e., the trajectory over time: orange solid line) is different from that with the optimal trait (orange dashed line) because of a time lag in adaptive evolution. **e-f**, Rapid evolution of species 2 results in coexistence with synchronous (**e**) or intermittent (**f**) cycles. Under intermittent cycles, the density of species 1 (blue solid line) increases when cycle amplitudes are small, whereas the density of species 2 (orange dashed line) increases when cycle amplitudes are large. The trait value of species 2 is depicted in black, and resource abundance in gray. Parameter values are *a*_1_ = 20, *α*_2_ = 84, *b*_1_ = *b*_2_ = 10, *d*_1_ = 0.8, *δ*_02_ = 1, *δ*_12_ = 0.2, *δ*_22_ = 4, *V*_1_ = 0, and *V*_2_ = 0.1 in **a**, **c**, and **e**, and *δ*_02_ = 0.25, *δ*_22_ = 12, and *V*_2_ = 0.08 in **b**, **d**, and **f**.

The underlying mechanism for coexistence (*sensu* Chesson 2000) can be understood by looking at the very-rapid-evolution limit. When evolution is much faster than ecological processes, the evolving species is always able to attain the optimal trait value that maximizes per-capita growth rate for a given resource concentration. The effect of very rapid evolution is to change the effective functional responses of the evolving species from concave down (gray curves in Fig. 2a-b) to concave up (especially when the limiting resource is scarce: orange curves in Fig. 2a-b) and it is this change that allows coexistence by enabling the evolving species to benefit from fluctuating environments in competition through nonlinear averaging. Although the realized per-capita growth rates when evolution is not much faster than ecology are different from those based on the optimal trait values due to the time lag in responses to changing environments (Fig. 2c-d, Fig. S4), coexistence is still possible because of relative nonlinearity.

It is important to note that rapid evolution not only broadens the parameter conditions that allow for coexistence via a so-called gleaner-opportunist trade-off (i.e., species 1 has a lower *R*\* whereas species 2 has a higher maximum growth rate (Grover 1990): Fig. 2a, c, and e), but it also yields stable coexistence under conditions previously thought to be incompatible with coexistence (Fig. 3). More specifically, coexistence is still possible even in the absence of a classical gleaner-opportunist trade-off, i.e., there is no trade-off between *R*\* and maximum growth rate, and the species with the smaller *R*\* and the higher maximum growth rate (species 1 in Fig. 2b, d, and f) is competitively impeded by fluctuations. Figure 3b shows that coexistence is not possible when genetic variance is zero, but as the genetic variance of the inferior species increases, the range of parameter values compatible with coexistence sharply increases. As such, this relaxation of the perceived restrictions characterizing classical relative nonlinearity can explain the coexistence of a much greater functional diversity of competing organisms than would be possible in the absence of rapid evolution.

**Figure 3.**
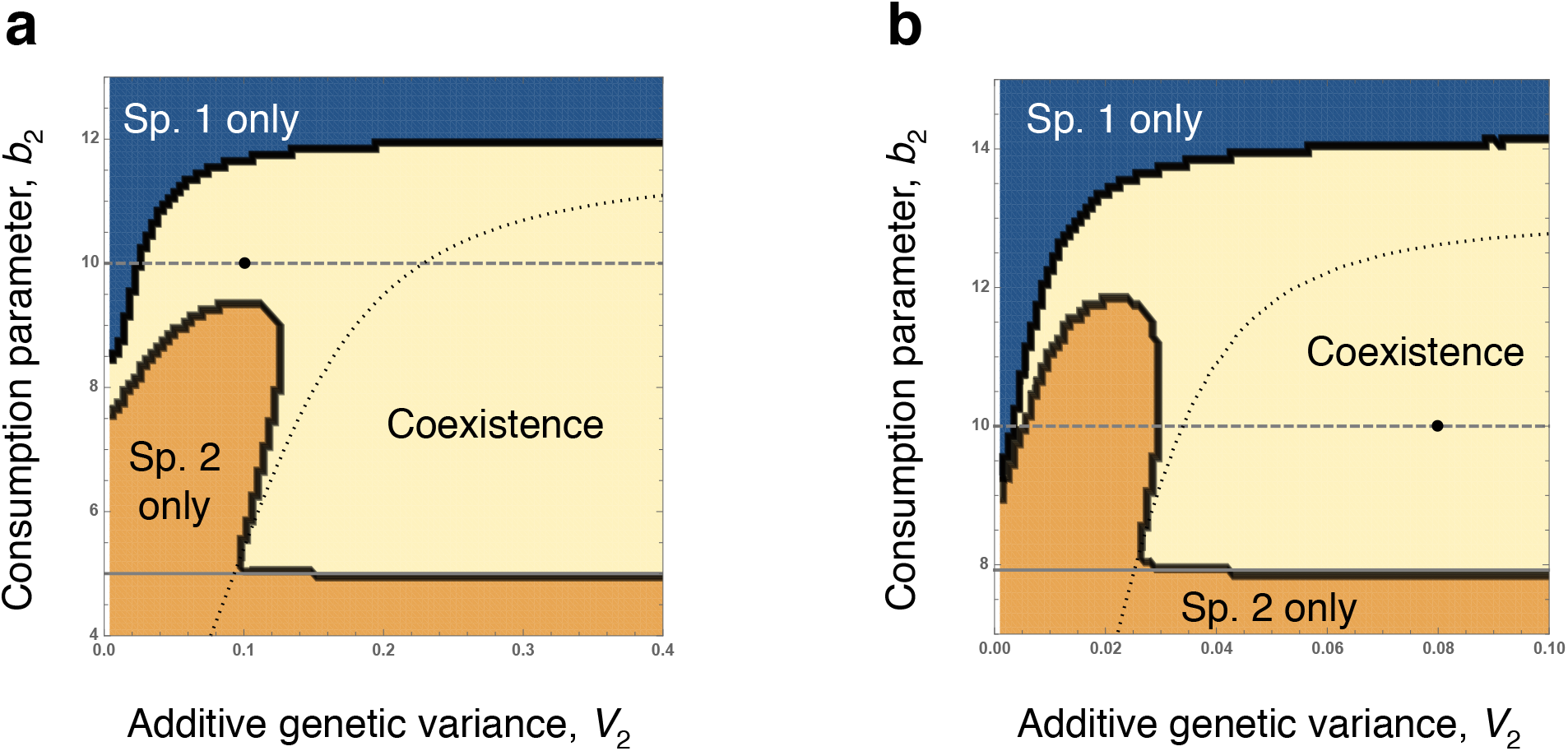
Effect of rapid evolution on parameter space yielding stable coexistence. **a-b**, Phase diagrams for the genetic variance, *V*_2_, and parameter that reduces the maximum consumption rate, *b*_2_, of species 2 where the per-capita growth rate of species 2 with the optimal trait is either larger than that of species 1 when the resource is abundant (**a**) or scarce (**b**). Dashed grey line indicates the situations where both species have the identical value of *b* (*b*_1_ = *b*_2_), where species 1 is competitively superior (lower *R*\*) and therefore excludes species 2 in the absence of genetic variance (Fig. 2, S2). At the limit of zero genetic variance, the coexistence region in **a**corresponds to classical relative nonlinearity. With increasing genetic variance, the parameter space corresponding to coexistence expands. Solid grey line indicates the point at which species 2 becomes the superior competitor and therefore genetic variance (in species 2) no longer promotes coexistence with smaller *b*_2_. Black dots indicate fixed parameter values in Figure 2. Black dotted curves indicate Hopf bifurcation in the system with species 2 and resource: the system is locally stable below the curves (with smaller *b*_2_ and larger *V*_2_).

## Discussion

Relative nonlinearity promotes species coexistence via fluctuations when (1) growth rates are relatively nonlinear functions of the limiting factor in competition (in this case, a shared resource) and (2) each species, when abundant, drives the amplitude of fluctuations in directions that favor their competitors (Fig. 1) (Yuan & Chesson 2015). Here we have shown that rapid evolution can satisfy the above two conditions under conditions previously thought to be incompatible with coexistence. Even without the classical gleaner-opportunist trade-off (Grover 1990), we show that coexistence is possible via relative nonlinearity as the evolving consumer species benefits from resource fluctuations in competition but stabilizes the consumer-resource cycles (Fig. 2–3).

Although our focus here is a system with rapidly evolving consumers competing for a single resource, a trade-off between competitive ability at equilibrium and the speed of adaptation may promote coexistence via relative nonlinearity under various other scenarios. For example, mechanisms of rapid adaptive trait changes include rapid evolution and phenotypic plasticity, and some organisms plastically suppress foraging activity and reproduction when resources are scarce (breeding suppression or dormancy), thereby stabilizing population dynamics (Nakazawa *et al.* 2009). This type of adaptive behavioral responses may also promote coexistence via relative nonlinearity (Appendix S4, Fig. S6). Recently, Tan *et al.* (2020) showed that a competitively inferior (larger *R*\*) consumer species with resource-dependent dormancy can coexist with a superior species in consumer-resource cycles. Although the authors discussed their findings in the context of game theory, our results indicate that they can also be explained as an outcome of relative nonlinearity. In addition, phenotypic plasticity in defense traits against predation can stabilize population dynamics (Yamamichi *et al.* 2019) and may also promote coexistence via relative nonlinearity. When the limiting factor in competition is predation pressure instead of resource abundance (i.e., apparent competition), species with higher predation tolerance (*P*\*) will persist (Holt 1977). However, coexistence is possible when species with lower predation tolerance stabilizes predator-prey cycles by inducible defenses (Appendix S5, Fig. S7). This provides an explanation for previous simulation results demonstrating coexistence of three prey types (asexually reproducing clones) with a shared predator and a shared resource (Yamamichi *et al.* 2011). According to classical *R*\* and *P*\* theory, only two species (or asexual clones) can coexist with a single predator and a single resource (Holt *et al.* 1994), but a third prey species with inducible defense can coexist with two non-plastic (defended and undefended) prey by relative nonlinearity. The destabilizing non-plastic prey have a competitive advantage when the dynamics are stable, while the stabilizing plastic prey have an advantage under fluctuations.

While our model assumptions are quite general, further theoretical studies are necessary to understand the generality and robustness of this novel mode of coexistence in ecosystems including, for example, the shape of the trade-off between consumption and mortality (Appendix S3, Fig. S5). Analyzing stochastic models will be particularly helpful to understand how low genetic variance (and therefore slower rates of evolution) in small populations impedes coexistence. It will be also valuable to conduct empirical studies that examine the effects of rapid evolution on species coexistence via relative nonlinearity. In addition to meta-analyses on the trade-off between foraging and mortality (Verdolin 2006) and the trade-off between competitive ability at equilibrium and the speed of adaptation, competition experiments will be valuable for deepening our understandings. Previous studies on the consequences of rapid evolution have typically used a single clone for competition experiments *after* long-term experimental evolution (Hart *et al.* 2019). However, it is rapid evolution itself during competition experiments that can promote coexistence via the mechanism outlined here, and a single clone cannot produce these kinds of dynamics. Although it is possible to manipulate the amount of genetic variation by changing the number of clones (Yoshida *et al.* 2003; Becks *et al.* 2010), we note that species with two clones will not coexist with another species via relative nonlinearity (mathematically, it is equivalent to three consumers competing for a single resource). In addition, one should note that rapid evolution does not always stabilize temporal fluctuations (Becks *et al.* 2010; Cortez & Patel 2017), in which case rapid evolution may result in priority effects via positive frequency-dependence in community dynamics (Ke & Letten 2018). Therefore, using phenotypically plastic species may be more feasible for showing coexistence driven by rapid adaptive trait dynamics via relative nonlinearity because phenotypic plasticity tends to strongly stabilize population dynamics, and the speed of adaptation will not be as slow as genetic evolution in small populations (Yamamichi *et al.* 2011; Yamamichi *et al.* 2019).

## Conclusion

With increasing evidence for rapid contemporary evolution on ecological timescales (Hendry 2016), an interest in developing a broader theory of species coexistence that incorporates rapid evolution and eco-evolutionary feedbacks has grown (Tachikawa 2008; Kremer & Klausmeier 2013; Wittmann & Fukami 2018; Hart *et al.* 2019; van Velzen 2020; Yamamichi *et al.* 2020). By allowing for rapid evolution in mechanistic resource competition models, we have identified a new solution to the paradox of the plankton, deriving from relative nonlinearity of competition. This mechanism may be important in competition between sexual and asexual species or species with different mutation rates. Alongside recent empirical work demonstrating relative nonlinearity across diverse systems (Letten *et al.* 2018; Hallett *et al.* 2019; Zepeda & Martorell 2019), we hope that our findings will help to motivate a reappraisal of the perceived importance of relative nonlinearity in nature.

## Acknowledgements

We thank S. P. Ellner, T. Fukami, and three anonymous reviewers for their helpful comments. MY thanks N. Shinohara for stimulating discussions during the initial stages of this project. MY was supported by the Japan Society for the Promotion of Science (JSPS) Grant-in-Aid for Scientific Research (KAKENHI) 18H02509 and 19K16223.

