## Supplementary Materials for "Rapid evolution promotes fluctuation-dependent species coexistence"

### Supplementary Material for "Rapid evolution promotes fluctuation-dependent species coexistence."

Last revision: December 11, 2020

**Contents**

Appendix S1: Necessary conditions for coexistence without evolution 2
Appendix S2: Consumer rapid evolution can stabilize fluctuations 5
Appendix S3: Trade-off between consumption and mortality 9
Appendix S4: Consumer phenotypic plasticity and intermittent cycles 10
Appendix S5: Prey inducible defense and intermittent cycles 12
References 13
Figure S1: Necessary conditions for coexistence without evolution 14
Figure S2: Competitive exclusion when there is no evolution 15
Figure S3: Local stability analyses when evolution is rapid 16
Figure S4: Functional responses and mortality rates of consumers in Fig. 1c, 2a, and 2b 18
Figure S5: Trade-off between consumption and mortality 19
Figure S6: Consumer phenotypic plasticity and intermittent cycles 20
Figure S7: Prey inducible defense and intermittent cycles 21

***Appendix S1: Necessary conditions for coexistence without evolution***

After nondimensionalization, Eq. 1 in the main text can be represented as follows:

$$\frac{dR'}{dt'} = R'\left[1 - R' - \sum_{j=1}^{2}\frac{a'_j\left(x_j\right)N'_j}{1+b'_jR'}\right],$$
$$\frac{dN'_i}{dt'} = N'_i\left[\frac{a'_i\left(x_i\right)R'}{1+b'_iR'} - d'_i\left(x_i\right)\right],\ \left(i=1,2\right), \qquad \text{(S1-1)}$$
$$\frac{dx_i}{dt'} = V_i\frac{\partial}{\partial x_i}\left(\frac{1}{N'_i}\frac{dN'_i}{dt'}\right),\ \left(i=1,2\right),$$

where $R' = R/K$, $N_i' = N_i/(c_iK)$, $t' = rt$, $a_i' = c_iKa_i/r$, $b_i' = b_iK$, and $d_i' = d_i/r$. We will omit the primes in the following. When $V_i = 0$, there are four necessary conditions for the two competing consumers to coexist (Hsu et al. 1978, Xiao and Fussmann 2013), but the coexistence region is determined by conditions 2-4 under our parameter settings. We plot the shaded region that satisfies the necessary conditions in the $d_1$-$d_2$ plane assuming that $b_1 > b_2$ (i.e., species 1 is competitively superior with lower $R^*$: Fig. S1a) and in the $b_1$-$b_2$ plane (Fig. S1b). The actual coexistence conditions are more restricted and can be checked by numerical simulations (Fig. S1). There are two types of coexistence: one is the case where there is a stable equilibrium of species 2 and the resource when species 1 is absent (Fig. S1c-d), and the other one is the case where species 2 shows limit cycles when species 1 is absent (Fig. S1e-f). When the two species have the same parameter value in $b$ (i.e., $b_1 = b_2$) as the main text, the following conditions 2 and 3 become $d_2 \le \frac{a_2d_1}{a_1}$ and $\frac{a_2d_1}{a_1} \le d_2$, respectively, and the two conditions cannot be satisfied unless $a_1d_2$ is exactly the same as $a_2d_1$.

1. $d_i < \frac{a_i}{1+b_i}$ (gray lines in Fig. S1a-b)

This implies that the equilibrium density of both consumers should be positive. When there is a single consumer and the other consumer is absent, the internal equilibrium is $\left(\bar{R},\bar{N}_i,\bar{N}_j\right)=\left(\frac{d_i}{a_i-b_id_i},\frac{a_i-\left(1+b_i\right)d_i}{\left(a_i-b_id_i\right)^2},0\right)$ ($i, j$ = 1, 2). As the denominator is squared, the consumer equilibrium density is positive when the numerator is positive.

2. $d_2 \le \frac{a_2\left(1+b_1\right)}{a_1\left(1+b_2\right)}d_1$ (orange lines in Fig. S1a-b)

This ensures that the competitively inferior species 2 (with higher $R^*$) has a higher growth rate than species 1 when the resource is abundant. When the resource level is at carrying capacity (i.e., $R$ = 1), species 2's mortality rate must be a smaller fraction of its per-capita growth rate compared to that of species 1: $\frac{d_2}{a_2/\left(1+b_2\right)-d_2} \le \frac{d_1}{a_1/\left(1+b_1\right)-d_1}$.

3. $\frac{a_2}{a_1/d_1-b_1+b_2} \le d_2$ (blue lines in Fig. S1a-b)

This means that species 1 is competitively superior in stable environments (i.e., species 1 has a lower $R^*$ than that of species 2). The $R^*$ is given by the equilibrium resource abundance in condition 1, and thus $\frac{d_1}{a_1-b_1d_1} \le \frac{d_2}{a_2-b_2d_2}$.

4. $1 < b_1$ and $d_1 < \frac{a_1\left(b_1-1\right)}{b_1\left(1+b_1\right)}$ (black lines in Fig. S1a-b)

As the competitively inferior species 2 cannot invade stable environments under condition 3 above, for coexistence the system with species 1 should show limit cycles (the equilibrium with resource and species 1 should be locally unstable). This can be obtained by checking the trace of the Jacobian matrix,

$$J = \left. \begin{pmatrix} \frac{\partial}{\partial R}\frac{dR}{dt} & \frac{\partial}{\partial N_1}\frac{dR}{dt} \\ \frac{\partial}{\partial R}\frac{dN_1}{dt} & \frac{\partial}{\partial N_1}\frac{dN_1}{dt} \end{pmatrix} \right|_{R=\bar{R},N_1=\bar{N}_1} = \begin{pmatrix} \frac{d_1\left[a_1(b_1-1)-b_1d_1(1+b_1)\right]}{a_1(a_1-b_1d_1)} & -d_1 \\ 1-\frac{(1+b_1)d_1}{a_1} & 0 \end{pmatrix}, \qquad \text{(S1-2)}$$

where the trace is $\frac{d_1\left[a_1(b_1-1)-b_1d_1(1+b_1)\right]}{a_1(a_1-b_1d_1)}$ and the determinant is $d_1\left[1-\frac{(1+b_1)d_1}{a_1}\right]$. The determinant is positive when condition 1 is satisfied, and thus the trace needs to be positive for the system to show limit cycles. As the denominator ($a_1 - b_1d_1$) is positive when there is an internal equilibrium ($\left(\bar{R},\bar{N}_1,\bar{N}_2\right)=\left(\frac{d_1}{a_1-b_1d_1},\frac{a_1-(1+b_1)d_1}{(a_1-b_1d_1)^2},0\right)$), the numerator determines the sign of the trace. This implies that the attacking rate should be large enough and the mortality should be small enough for the system to exhibit limit cycles.

Under the parameter settings of Fig. 2 in the main text, coexistence is impossible as the two consumers have the same parameter value in $b$ ($b_1 = b_2$) and there is no parameter combination that satisfies the conditions 2 and 3 (Fig. S2a-f). In this case, coexistence is possible only when evolution is rapid enough (Fig. S2g-h).

***Appendix S2: Consumer rapid evolution can stabilize fluctuations***

As previous studies showed that consumer rapid evolution can cause or stabilize limit cycles depending on model assumptions (Cortez and Patel 2017), here we show that rapid evolution stabilizes limit cycles in our model. We consider a system with a single evolving consumer and resource:

$$\begin{aligned}\frac{dR}{dt} &= R\left[1 - R - \frac{a(x)N}{1+bR}\right],\\ \frac{dN}{dt} &= N\left[\frac{a(x)R}{1+bR} - d(x)\right],\\ \frac{dx}{dt} &= V\frac{\partial}{\partial x}\left(\frac{1}{N}\frac{dN}{dt}\right),\end{aligned} \qquad \text{(S2-1)}$$

where $a(x) = \alpha x$ and $d(x) = \delta_0 + \delta_1 x + \delta_2 x^2$, assuming that $a$ and $d$ are increasing functions of $x$. The coefficient for the attacking rate, $\alpha$, the intercept and coefficients for the mortality, $\delta_0$, $\delta_1$, and $\delta_2$, and the additive genetic variance, $V$, are assumed to be positive. The trait dynamics can be written as:

$$\frac{dx}{dt} = V\left(\frac{\alpha R}{1+bR} - \delta_1 - 2\delta_2 x\right), \qquad \text{(S2-2)}$$

When there is no evolution ($V = 0$), the system shows limit cycles when $1 < b$ and $d < \frac{a(b-1)}{b(1+b)}$ (Appendix S1, Fig. S3a-c). With evolution ($V > 0$), the system is locally stable when the real parts of the eigenvalues of the following Jacobian matrix are negative:

$$J=\begin{pmatrix} J_{11} & J_{12} & J_{13} \\ J_{21} & J_{22} & J_{23} \\ J_{31} & J_{32} & J_{33} \end{pmatrix} = \left.\begin{pmatrix} \frac{\partial}{\partial R}\frac{dR}{dt} & \frac{\partial}{\partial N}\frac{dR}{dt} & \frac{\partial}{\partial x}\frac{dR}{dt} \\ \frac{\partial}{\partial R}\frac{dN}{dt} & \frac{\partial}{\partial N}\frac{dN}{dt} & \frac{\partial}{\partial x}\frac{dN}{dt} \\ \frac{\partial}{\partial R}\frac{dx}{dt} & \frac{\partial}{\partial N}\frac{dx}{dt} & \frac{\partial}{\partial x}\frac{dx}{dt} \end{pmatrix}\right|_{R=R^*,N=N^*,x=x^*} . \qquad \text{(S2-3)}$$

The sign structure of the Jacobian matrix is

$$\begin{pmatrix} \pm & - & - \\ + & 0 & 0 \\ \pm & 0 & \pm \end{pmatrix}, \qquad \text{(S2-4)}$$

where ± denotes the entries that can have either sign. Because of the consumption interaction, increasing the consumer density reduces the resource abundance ($J_{12} < 0$), whereas increasing the resource abundance increases the consumer density ($J_{21} > 0$). Increasing the consumer trait reduces the resource abundance ($J_{13} < 0$). $J_{23}$ is zero because we assume frequency independent selection (i.e., fitness is solely determined by an individual's trait value and not affected by the population mean trait value). We assume that the determinant of the Jacobian is negative ($|J| < 0$) so that equilibria and cycles arising via Hopf bifurcations are attractors (Cortez and Patel 2017). As $|J| = -J_{12}J_{21}J_{33}$ in our model, $J_{33} < 0$ when the determinant is negative. This means that evolution is stabilizing rather than disruptive, and increasing the additive genetic variance can stabilize dynamics (Cortez and Patel 2017). In addition, this implies $J_{31} > 0$ in our model as $J_{31} = CJ_{33}$ where $C$ is a negative constant.

More specifically, local stability of internal equilibrium can be examined by the Routh-Hurwitz criteria based on the characteristic polynomial of the Jacobian matrix,

$\rho(\lambda) = \lambda^3 + a_1\lambda^2 + a_2\lambda + a_3$, where

$$\begin{aligned} a_1 &= -\left(J_{11} + VJ_{33}\right), \\ a_2 &= V\left(J_{11}J_{33} - J_{13}J_{31}\right) - J_{12}J_{21}, \\ a_3 &= VJ_{12}J_{21}J_{33}. \end{aligned} \tag{S2-5}$$

Here we change the notation from $J_{3i}$ to $VJ_{3i}$ ($i = 1, 3$) so that readers can see the effects of additive genetic variance, $V$, on local stability. The equilibrium undergoes a Hopf bifurcation when $0 < a_1$ and $a_1a_2 - a_3 = 0$. As evolutionary dynamics is stable ($J_{33} < 0$), we consider the following two cases: (1) when ecological dynamics is stable, and (2) when ecological dynamics is unstable.

When ecological dynamics is stable ($-J_{12}J_{21} > 0$ and $J_{11} < 0$), the system is stable when $V$ is small. In addition, $a_1$ is positive and each coefficient of the following equation,

$$a_1a_2 - a_3 = \underbrace{J_{33}\left(J_{13}J_{31} - J_{11}J_{33}\right)}_{\text{positive}}V^2 + \underbrace{J_{11}\left(J_{13}J_{31} - J_{11}J_{33}\right)}_{\text{positive}}V + \underbrace{J_{11}J_{12}J_{21}}_{\text{positive}}, \tag{S2-6}$$

is positive. Therefore, there is no Hopf bifurcation and the system is locally stable irrespective of $V$ values.

When ecological dynamics is unstable ($-J_{12}J_{21} > 0$ and $J_{11} > 0$), on the other hand, the system is unstable with sufficiently small $V$ values. When $J_{11}J_{33} - J_{13}J_{31} > 0$, $a_1a_2$ is negative when $V = 0$, and has a positive linear coefficient and a negative quadratic coefficient. This means that $a_1a_2$ is equal to $a_3$ with two $V$ values or a single $V$ value, or $a_1a_2$ is always smaller than $a_3$. In the third case, the equilibrium is unstable for all $V$. In the second case, the equilibrium becomes neutrally stable at the $V$ value and unstable otherwise. In the first case, a

Hopf bifurcation can occur at the two $V$ values, and the equilibrium is locally stable for intermediate values of $V$ and unstable otherwise. When $J_{11}J_{33} - J_{13}J_{31} < 0$, $a_1a_2$ has a positive quadratic coefficient and thus there is a single value of $V$ where $a_1a_2 - a_3 = 0$. This means that the equilibrium is unstable for smaller values of $V$ and stable for larger values of $V$. In our parameter setting, $J_{11}J_{33} - J_{13}J_{31} < 0$ and thus rapid evolution stabilizes fluctuations (Fig. S3d-f).

In addition to the analyses based on the 3×3 Jacobian matrix (Eq. S2-3), we can obtain further insights by considering the situation where evolutionary dynamics is much faster than ecological dynamics ($V \to \infty$). The optimal trait at a certain resource level, $R^*$, is given by

$$x^* = \frac{1}{2\delta_2}\left(\frac{\alpha R^*}{1+bR^*} - \delta_1\right). \qquad \text{(S2-7)}$$

In this case, we can check the local stability of the 2×2 Jacobian matrix as Eq. S1-2. The determinant is positive when $\alpha > (1+b_1)\left(\delta_1 + 2\sqrt{\delta_0\delta_2}\right)$ and the trace is negative when $\alpha^2 + \delta_1 b_1(1+b_1)\left(\delta_1 + 6\sqrt{\delta_0\delta_2}\right) > \alpha\left(\delta_1 + 2\delta_1 b_1 + 6b_1\sqrt{\delta_0\delta_2}\right)$. This implies that the system is locally stable when the consumption parameter, $b_1$, mortality intercept, $\delta_0$, and mortality quadratic coefficient, $\delta_2$, are small (Fig. S3g-i).

***Appendix S3: Trade-off between consumption and mortality***

We assume that a quantitative trait $x$ affects the consumption rate parameter $a$ linearly and the mortality rate $d$ quadratically in the main text. According to Appendix S2, $d$ needs to be a convex function of $x$ as long as $a$ is a linear function of $x$ and the functional response is $aR/(1+hR)$. In our model, $J_{33} < 0$ is necessary for the determinant of the Jacobian to be negative and equilibria and cycles arising via Hopf bifurcations are attractors (Cortez and Patel 2017). Because $J_{33} = \partial(dx/dt)/\partial x = -V\partial^2 d/\partial x^2$, $J_{33} < 0$ implies $\partial^2 d/\partial x^2 > 0$ (i.e., $d$ is a convex function of $x$).

Then, how does the exponent of the convex function affect coexistence? To simplify the notation, we regard the parameter $a$ as an evolving variable hereafter. Thus, the mortality rate $d$ is a function of $a$ (e.g., $d(a) = d_0 + d_1 a + d_2 a^2$). When the mortality rate is an increasing function of the consumption parameter (i.e., $\partial d/\partial a > 0$), there are three possibilities: (1) the mortality rate is a concave function of $a$ (i.e., $\partial^2 d/\partial a^2 < 0$), (2) the mortality rate is a linear function of $a$ (i.e., $\partial^2 d/\partial a^2 = 0$), and (3) the mortality rate is a convex function of $a$ (i.e., $\partial^2 d/\partial a^2 > 0$). As the mortality rate needs to be a convex function of $a$ (3) for equilibria and cycles arising via Hopf bifurcations to be attractors (Cortez and Patel 2017), we examined how the exponent affects coexistence (Fig. S5). We found that large exponents (i.e., more convex functions) result in the loss of trade-off as the evolving consumer can increase its consumption rate while keeping its mortality rate small (i.e., the consumer can approach the bottom-right region in Fig. S5a). Therefore, intermediate exponent values are necessary for coexistence with rapid evolution via relative nonlinearity (Fig. S5).

***Appendix S4: Consumer phenotypic plasticity and intermittent cycles***

Here we show that a plastic consumer with resource-dependent dormancy can show the same intermittent cycles as the evolution model in the main text. We assume that the plastic consumer changes its trait in response to resource abundance (Tan et al. 2020):

$$\begin{aligned}
\frac{dR}{dt} &= R\left(1 - R - \frac{a_1 N_1}{1+b_1 R} - \frac{a_2 N_{21}}{1+b_2 R}\right),\\
\frac{dN_1}{dt} &= N_1\left(\frac{a_1 R}{1+b_1 R} - d_1\right),\\
\frac{dN_{21}}{dt} &= N_{21}\left(\frac{a_2 R}{1+b_2 R} - d_{21}\right) - f_1(R)N_{21} + f_2(R)N_{22},\\
\frac{dN_{22}}{dt} &= f_1(R)N_{21} - f_2(R)N_{22} - d_{22}N_{22},
\end{aligned} \qquad \text{(S4-1)}$$

where the active consumer, $N_{21}$, has a higher mortality rate than the dormant consumer, $N_{22}$: $d_{21} > d_{22}$. We assume the following reaction norm functions,

$$\begin{aligned}
f_1(R) &= \frac{1}{1+\left(R/g\right)^b},\\
f_2(R) &= 1 - f_1(R),
\end{aligned} \qquad \text{(S4-2)}$$

where $b$ is the resource threshold and $g$ is the sensitivity to resource abundance (Vos et al. 2004), and thus $f_1(R)$ is a decreasing function of the resource abundance and $f_2(R)$ is an increasing function of the resource abundance (Fig. S6a). Here we again assumed $b_1 = b_2$ and the non-plastic species (species 1) has the lowest resource requirement ($R^*$). When there is no plasticity ($f_1 = f_2 = 0$), the model (Eq. S4-1) is equivalent to the case with two consumers competing for a single resource (as $N_{22}$ decreases to zero when $f_1 = f_2 = 0$). Coexistence via relative nonlinearity is impossible because there is no parameter combination that satisfies the conditions 2 and 3 of Appendix S1: the species 1 has the lowest $R^*$, and the other consumer's growth rate is lower than that of species 1 even when the resource is abundant. Hence, competitive exclusion occurs without plasticity (Fig. S6b). However, coexistence becomes possible with phenotypic plasticity, resulting in intermittent cycles (Fig. S6c). When the fraction of the active consumer is $N_{21}/(N_{21} + N_{22}) = f_2(R)$ and that of the dormant consumer is $N_{22}/(N_{21} + N_{22}) = f_1(R)$, the per-capita growth rate is:

$$\frac{1}{N_2}\frac{dN_2}{dt} = f_2(R)\left(\frac{a_2 R}{1+b_2 R} - d_{21}\right) - f_1(R) d_{22}, \qquad \text{(S4-3)}$$

and this results in a convex function when the resource is scarce (relative nonlinearity: Fig. S6d).

***Appendix S5: Prey inducible defense and intermittent cycles***

We show that prey inducible defense can also result in species coexistence. Here the predator density, $P$, and the two prey species densities, $N_1$ and $N_2$, are

$$\begin{aligned}\frac{dP}{dt} &= P\left(\frac{a_1N_1 + a_2N_{21}}{1+b_1N_1+b_2N_{21}} - d_1\right),\\ \frac{dN_1}{dt} &= N_1\left[r_1\left(1-N_1\right) - \frac{a_1P}{1+b_1N_1+b_2N_{21}}\right],\\ \frac{dN_{21}}{dt} &= N_{21}\left[r_2\left(1-N_{21}-N_{22}\right) - \frac{a_2P}{1+b_1N_1+b_2N_{21}}\right] - f_2(P)N_{21} + f_1(P)N_{22},\\ \frac{dN_{22}}{dt} &= f_2(P)N_{21} - f_1(P)N_{22} - d_2N_{22},\end{aligned} \tag{S5-1}$$

where $N_{12}$ is the active prey density and $N_{22}$ is the defended prey density ($N_2 = N_{21} + N_{22}$). The reaction norm function is the same as Eq. S4-2, and hence $f_1(P)$ is a decreasing function of the predator density and $f_2(P)$ is an increasing function of the predator density (Fig. S7a). Here we assumed $a_1 = a_2$, $b_1 = b_2$, $r_1 > r_2$, and the non-plastic species (species 1) has the larger predator tolerance ($P^*$). When there is no plasticity ($f_1 = f_2 = 0$), the model (Eq. S5-1) is equivalent to the case with two prey species with a predator (as $N_{22}$ decreases to zero when $f_1 = f_2 = 0$). Coexistence does not occur because of the difference in prey growth ($r_1 > r_2$) and the equivalent predation pressures ($a_1 = a_2$ and $b_1 = b_2$: Fig. S7b). However, coexistence becomes possible with phenotypic plasticity in intermittent cycles (Fig. S7c). When the fraction of the active prey is $N_{21}/(N_{21} + N_{22}) = f_1(P)$ and that of the defended prey is $N_{22}/(N_{21} + N_{22}) = f_2(P)$, the per-capita growth rate is in a convex function when the predator is scarce (relative nonlinearity: Fig. S7d).

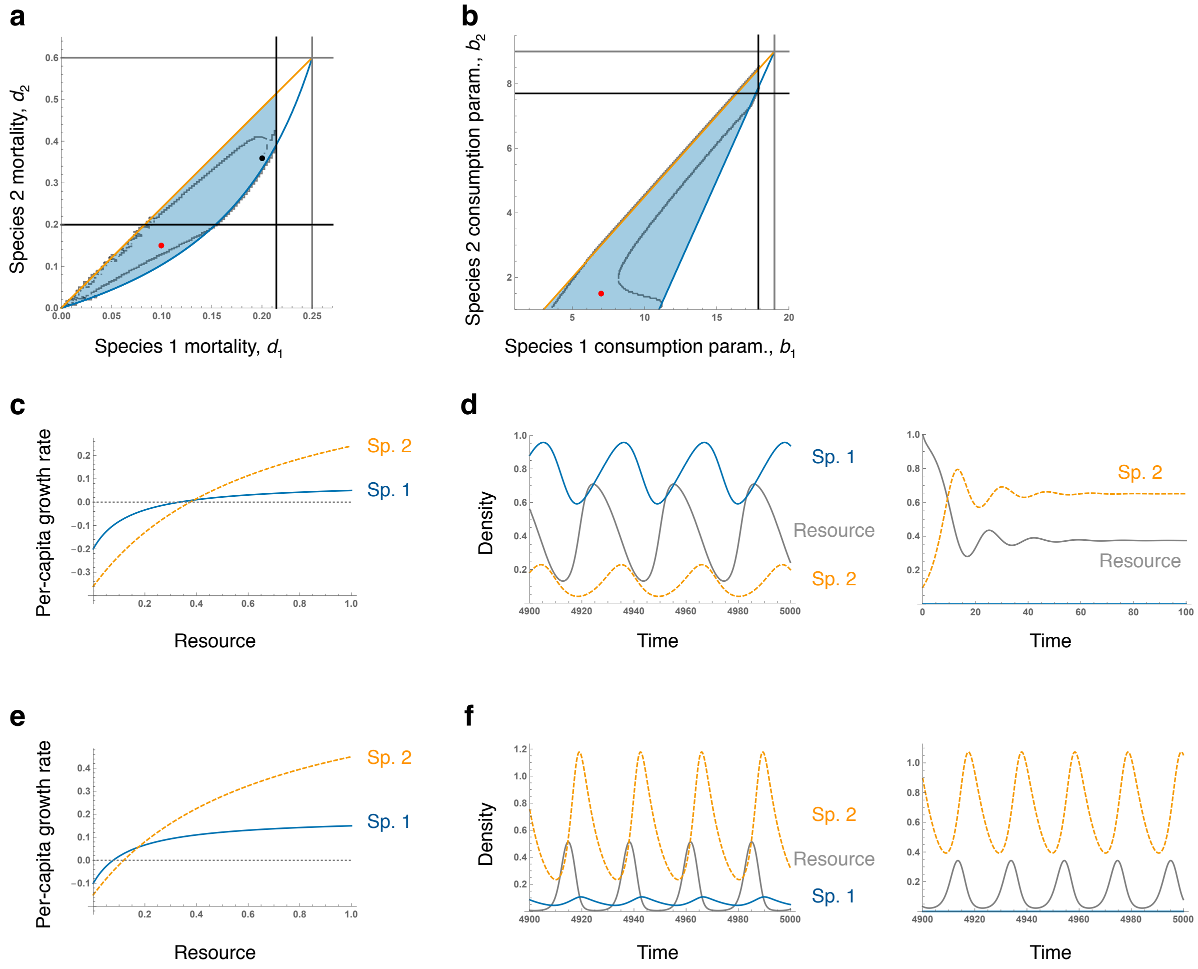


**Figure S1 | Necessary conditions for coexistence without evolution. a**, A phase diagram for four necessary conditions for coexistence with mortality rates, $d$. The gray lines indicate condition 1, the orange line indicates condition 2, the blue curve indicates condition 3, and the black lines indicate condition 4 (Hopf bifurcations). Numerical simulations confirmed that coexistence is possible in the region surrounded by black points. Parameter values are $a_1 = 2$, $a_2 = 1.5$, $b_1 = 7$, and $b_2 = 1.5$. **b**, A phase diagram with consumption parameters, $b$. Parameter values are $a_1 = 2$, $a_2 = 1.5$, $d_1 = 0.1$, and $d_2 = 0.15$. **c-d**, Species coexistence when $d_1 = 0.2$ and $d_2 = 0.36$ (a black point in **a**). Species 2 exhibits stable dynamics in the absence of species 1. **e-f**, Species coexistence when $d_1 = 0.1$ and $d_2 = 0.15$ (red points in **a** and **b**). Species 2 exhibits limit cycles in the absence of species 1.

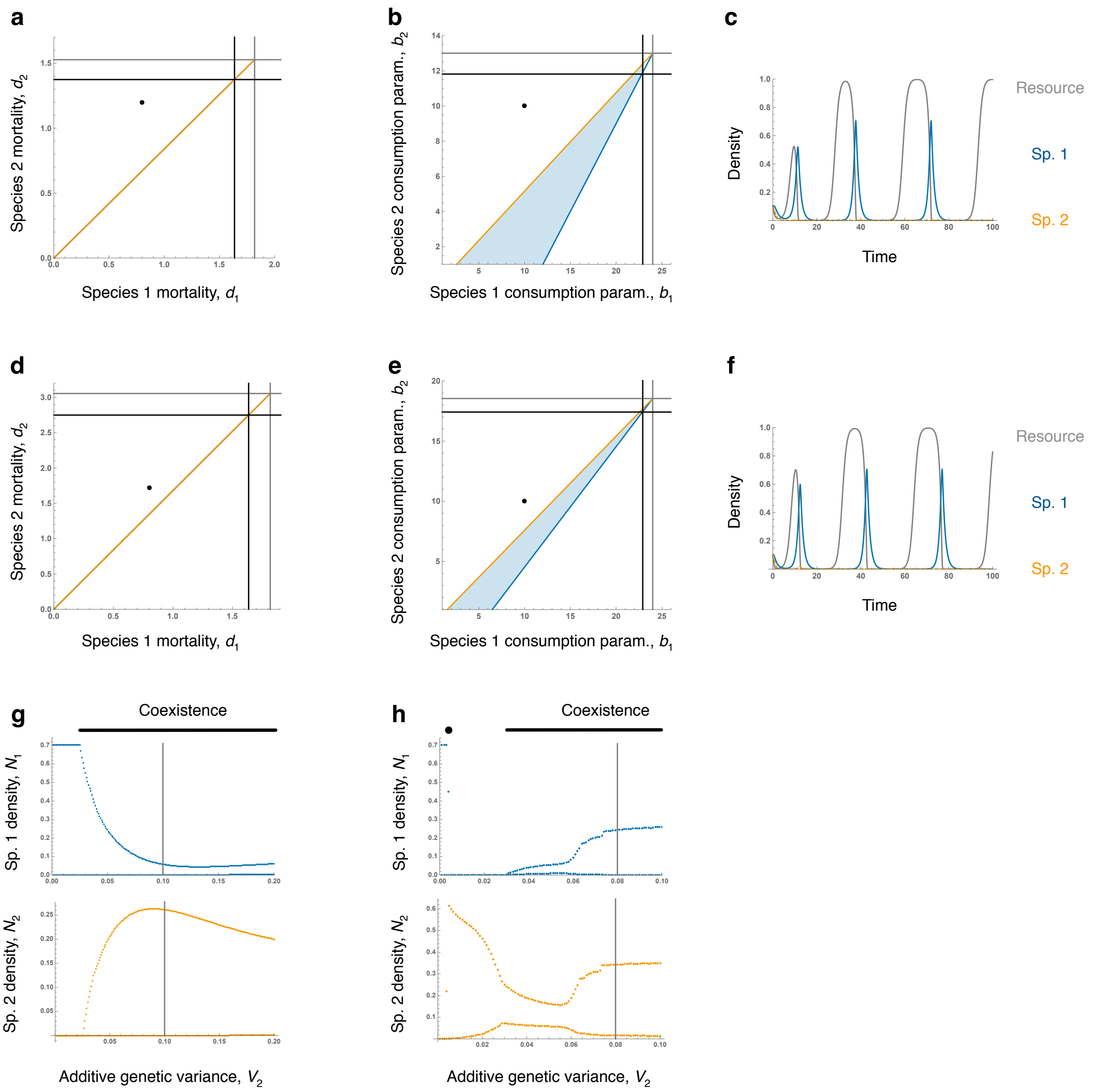


**Figure S2 | Competitive exclusion when there is no evolution. a-c**, When the trait value ($x_2$) is 0.2 under the parameter settings of Fig. 2a, c, and e in the main text. Because the two consumers have the same consumption rate parameter ($b_1 = b_2$), coexistence is not possible (black points in the phase diagrams of **a** and **b**), and consumer 2 goes extinct (**c**). **d-f**, When the trait value ($x_2$) is 0.4. **g-h**, Bifurcation plots of the additive genetic variance of consumer 2, $V_2$, show that the two consumers coexist when evolution is rapid under the parameter settings of Fig. 2a, c, and e (**g**) and Fig. 2b, d, and f (**h**). Gray lines show the parameter condition in Fig. 2 ($V_2 = 0.1$ or $0.08$).

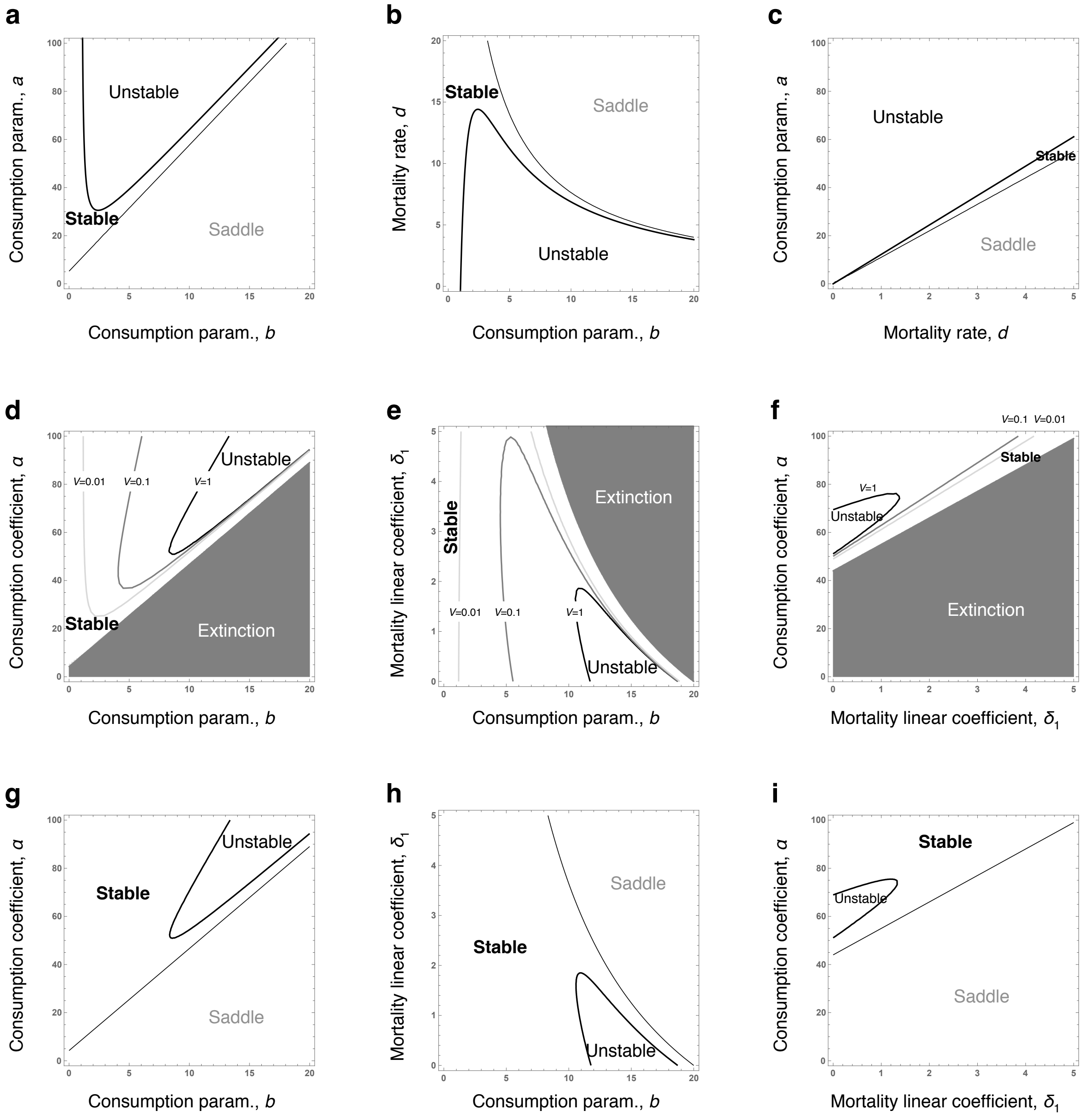


**Figure S3 | Local stability analyses with no evolution (a-c), rapid evolution (d-f), and very rapid evolution limit (g-i). a-c**, Phase diagrams for the consumption rate parameters, *b* and *a*, and mortality rate, *d*, when there is no evolution. Local stability analyses based on the condition 4 in Appendix S1 are shown. Stable, unstable, and saddle regions are for locally stable equilibria, locally unstable equilibria (with limit cycles), and saddle points, respectively. **d-i**, Phase diagrams for the consumption rate parameter, *b*, consumption rate coefficient, $\alpha$, and mortality linear coefficient, $\delta_1$, with rapid evolution. Local stability analyses based on the 3×3 Jacobian matrix (Eq. S2-3) with $V = 0.01$, 0.1, and 1 (**d-f**) as well as the very rapid evolution limit (**g-i**) are shown. Basic parameter values are $a = 84$, $b = 10$, and $d = 5.24$ in **a-c**, and $\alpha = 84$, $b = 10$, $\delta_0 = 1$, $\delta_1 = 0.24$, and $\delta_2 = 4$ in **d-i**.

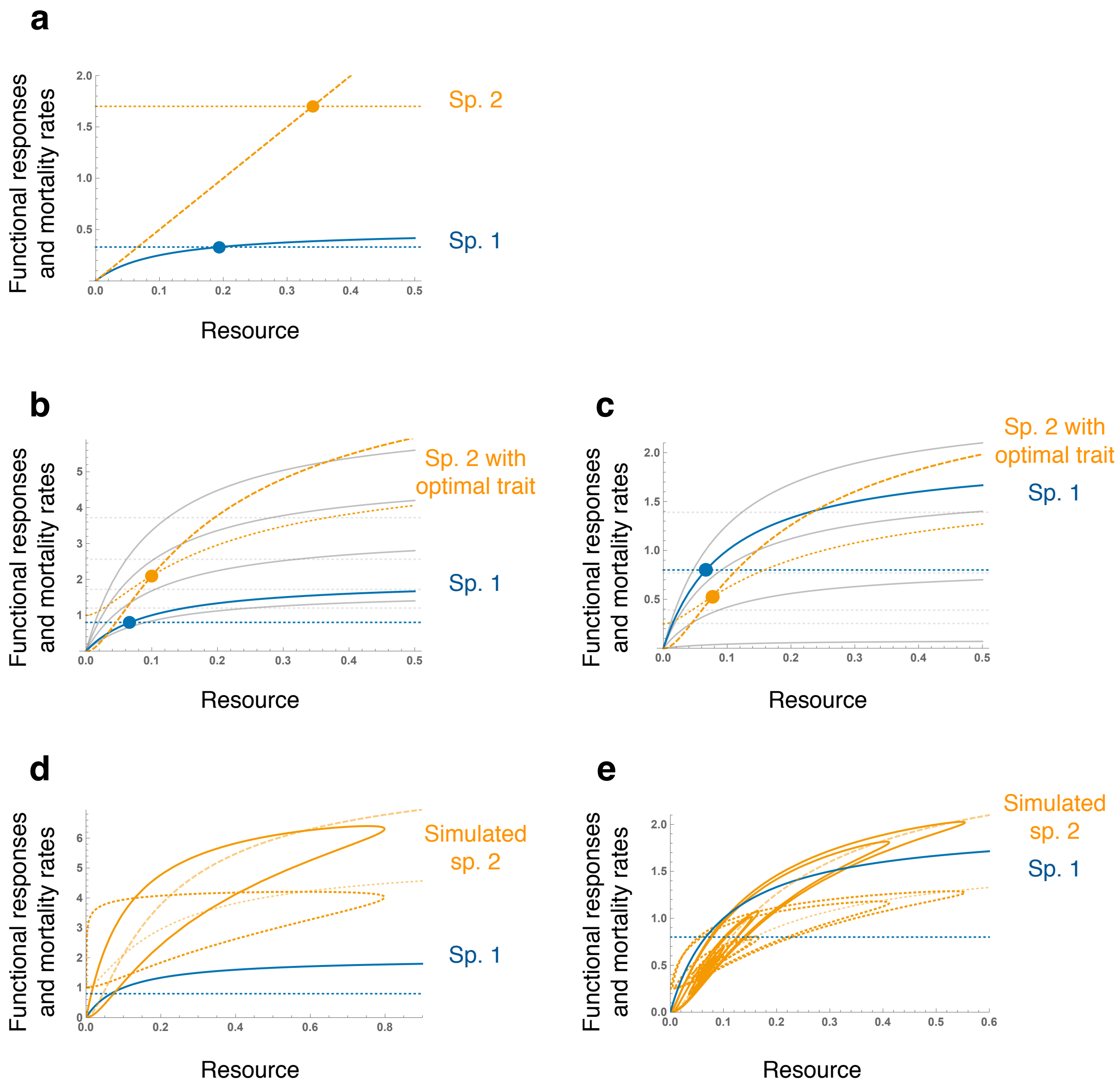


**Figure S4 | Functional responses and mortality rates of consumers in Fig. 1c, 2a, and 2b.** **a**, Functional responses and mortality rates of species 1 (blue) and 2 (orange) in Fig. 1c in the main text. **b-e**, Functional responses and mortality rates of non-evolving species 1 (blue) and evolving species 2 (orange) in Fig. 2a (**b**, **d**) and Fig. 2b (**c**, **e**). Functional responses and mortality rates with the optimum trait (**b-c**) and the simulated trait trajectory (**d-e**) are shown.

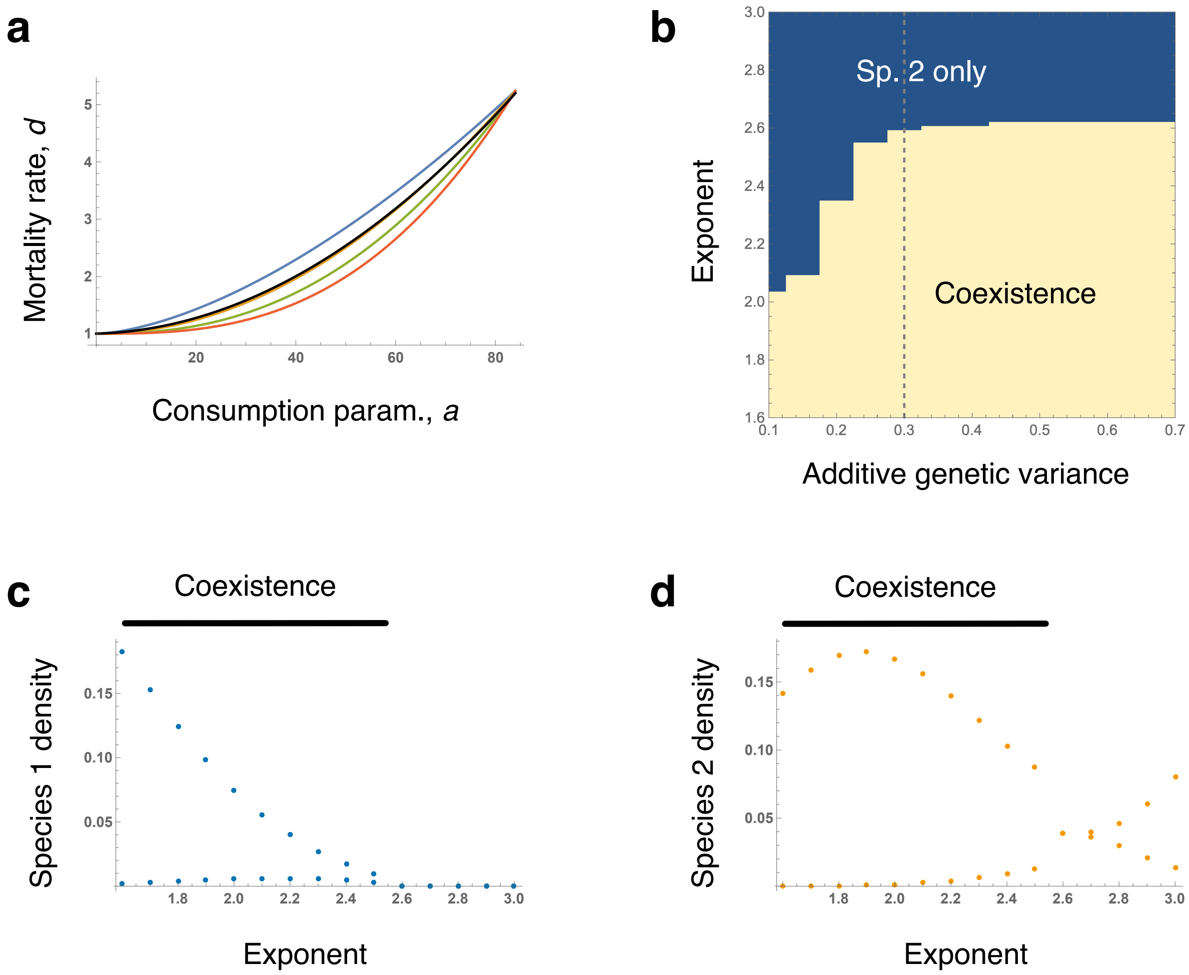


**Figure S5 | Trade-off between consumption and mortality. a**, The trade-off between the consumption rate parameter, $a$, and mortality rate, $d$. The black curve is the function in Fig. 2a, c, e, and 3a. Other curves show trade-off with different exponents: $d = d_0(a/a_0)^\alpha + d_1$, and $\alpha = 1.6$ (blue), 2 (orange), 2.4 (green), and 2.8 (red). Other parameter values are $d_0 = 4.24$, $a_0 = 84$, and $d_1 = 1$. **b**, A phase diagram showing the effects of consumer additive genetic variance, $V_2$, and exponent, $\alpha$, on coexistence. Dashed gray line indicates parameter conditions for **c-d**. **c-d**, Bifurcation plots showing the effects of the exponent on population dynamics of non-evolving species 1 (**c**) and evolving species 2 (**d**).

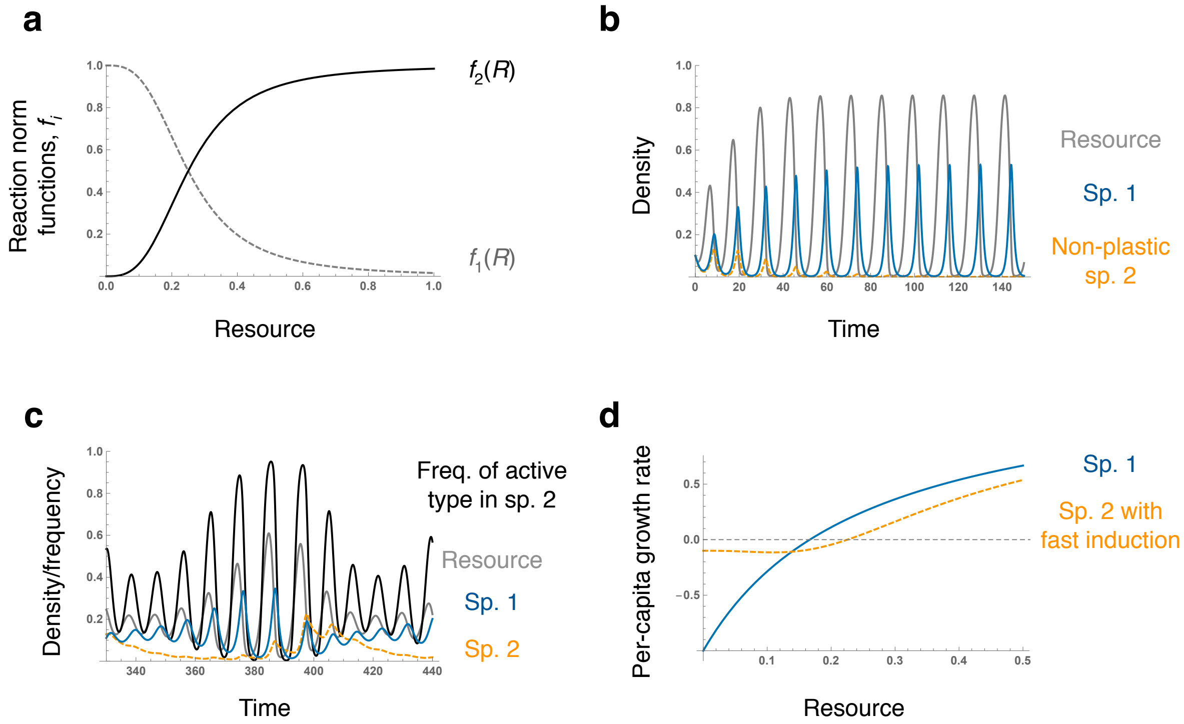


**Figure S6 | Consumer phenotypic plasticity and intermittent cycles. a**, Consumer reaction norm functions (Eq. S4-2) with $g = 0.25$ and $b = 3$. **b**, When consumer 2 (orange dashed line) has no plasticity, coexistence does not occur and consumer 1 (blue solid line) persists with resource (gray). **c**, Phenotypic plasticity in consumers causes intermittent cycles. The frequency of the active type in species 2 (black) increases when resource is abundant. This results in intermittent cycles where the plastic consumer increases in fluctuating environments whereas the non-plastic consumer increases in stable environments. **d**, When the plastic response is fast, the plastic consumer shows convex down per-capita growth rate whereas the non-plastic consumer has lower $R^*$. Parameter values are $a_1 = a_2 = 10$, $b_1 = b_2 = 4$, $d_1 = 1$, $d_{21} = 1.05$, and $d_{22} = 0.1$.

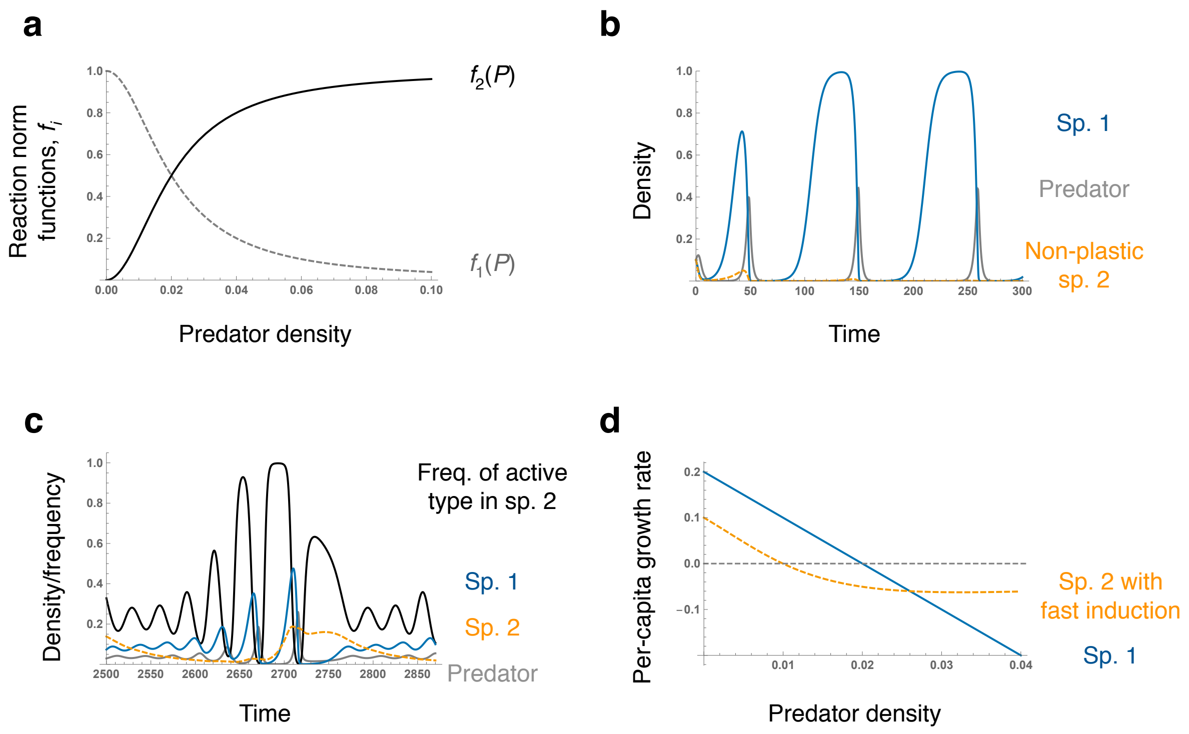


**Figure S7 | Prey inducible defense and intermittent cycles.** **a**, Prey reaction norm functions with $g = 0.02$ and $b = 2$. **b**, When prey species 2 (orange dashed line) has no plasticity, coexistence does not occur and the predator (gray line) and prey species 1 (blue solid line) persist. **c**, Prey inducible defense promotes species coexistence in intermittent cycles. The frequency of active prey in species 2 (black) increases when predator is scarce. This results in intermittent cycles where the plastic prey increases in fluctuating environments whereas the non-plastic prey increases in stable environments. **d**, When the plastic response is fast, the plastic prey shows convex down per-capita growth rate whereas the non-plastic prey has higher $P^*$ when rare ($N_1 = N_2 = 0$). Parameter values are $a_1 = a_2 = 10$, $b_1 = b_2 = 10$, $d_1 = 0.5$, $d_2 = 0.001$, $r_1 = 0.2$, and $r_2 = 0.1$.
